# Hemodynamic phenotypes linked to high-altitude subclinical organ damage

**DOI:** 10.64898/2026.04.17.719322

**Authors:** Chao Huijuan, Bao Guoping, Wang Xuewei, Tang Biwen, Wang Qian, Hu Yueliang, Alberto Avolio, Zuo Junli

**Affiliations:** Department of Geriatrics, Ruijin Hospital, Shanghai Jiao Tong University School of Medicine, Shanghai, China; Department of Cardiology Deqing Tibetan Autonomous Prefecture People’s Hospital; Zhenxin Community Health Service Center, Jiading District, Shanghai; Faculty of Medicine, Health and Human Sciences, Talavera Road, Macquarie University, Sydney, NSW 2109, Australia

**Keywords:** High altitude, Hemodynamic phenotypes, Vascular stress, Arterial stiffness, Target organ damage, Cluster analysis

## Abstract

**Background:** Chronic exposure to high-altitude hypoxia imposes sustained cardiovascular stress, yet hemodynamic adaptation among healthy high-altitude dwellers is heterogeneous and remains poorly characterized. This study aimed to identify distinct hemodynamic phenotypes in a healthy high-altitude population using unsupervised machine learning and to evaluate their association with multi-system subclinical target organ damage.

**Methods:** This cross-sectional study enrolled 694 healthy adults permanently residing at ≥3300 m on the Qinghai-Tibet Plateau. Unsupervised K-means clustering was performed on nine hemodynamic variables, including peripheral and central blood pressures, augmentation index (AIx), pulse pressure amplification ratio (pPP/cPP), and systolic pressure amplification (pSBP-cSBP). Differences across phenotypes in carotid intima-media thickness (IMT), estimated glomerular filtration rate (eGFR), left ventricular mass index (LVMI), and pulse wave velocity (PWV) were assessed using one-way ANOVA with Bonferroni-corrected post-hoc tests.

**Results:** Three distinct hemodynamic phenotypes were successfully identified. The **C2 (Balanced Adaptation)** phenotype (n = 245) demonstrated the most favorable hemodynamic profile, characterized by the lowest blood pressure and augmentation index (AIx) values, along with the highest peripheral-to-central pulse pressure ratio (pPP/cPP). The **C1 (Vascular Stress)** phenotype (n = 267) presented with normal peripheral systolic blood pressure (125.9 ± 11.3 mmHg) but exhibited markedly elevated wave reflection indices, including the highest heart rate-adjusted augmentation index (AIx@HR75: 31.9 ± 9.7%) and the lowest pPP/cPP ratio (1.29 ± 0.08). The **C3 (High-Load Decompensation)** phenotype (n = 182) displayed significantly elevated blood pressures and the greatest overall hemodynamic load. Regarding target organ damage, a clear gradient was observed across the three phenotypes. The C3 phenotype showed the highest carotid intima-media thickness (IMT: 1.162 ± 0.23 mm) and left ventricular mass index (LVMI: 69.18 ± 40.73 g/m²). Conversely, the C2 phenotype exhibited the highest estimated glomerular filtration rate (eGFR: 97.38 ± 16.38 mL/min/1.73m²) and the lowest IMT (0.994 ± 0.26 mm). The C1 phenotype consistently displayed intermediate values for all organ damage indicators. After Bonferroni correction, all pairwise comparisons for LVMI and pulse wave velocity (PWV) reached statistical significance (all P < 0.05).

**Conclusions:** Healthy high-altitude individuals manifest three distinct hemodynamic phenotypes arrayed along a cardiovascular risk continuum. The novel Vascular Stress (C1) phenotype represents a “masked” high-risk state characterized by normal peripheral blood pressure but elevated arterial stiffness and wave reflection, challenging sole reliance on brachial pressure for risk assessment. This phenotype-based stratification provides a framework for precision prevention and early intervention in high-altitude populations.

## 1 Introduction

Chronic exposure to the hypobaric-hypoxic environment at high altitude presents a persistent physiological challenge to the human cardiovascular system. The body initiates a suite of compensatory hemodynamic adjustments, including increased heart rate, elevated pulmonary artery pressure, and alterations in peripheral vascular resistance, to maintain adequate tissue oxygen delivery^1^. While these mechanisms support short-term survival, their long-term consequences may paradoxically contribute to increased cardiovascular stress and accelerate the development of subclinical organ damage. Notably, markers of vascular dysfunction and early target organ injury, such as increased arterial stiffness and elevated left ventricular mass, are established independent predictors of future major adverse cardiovascular events ^2^. Understanding the drivers of this preclinical damage in high-altitude populations is therefore a public health priority ^3^.

However, current research paradigms in high-altitude medicine are limited by a focus on isolated physiological metrics (e.g., blood pressure, PWV) ^4^. Numerous studies have documented group-level differences in these specific parameters between highland and lowland populations^5,6^. While invaluable, this approach treats the ostensibly “healthy” high-altitude population as a homogeneous entity, analyzing variables in isolation. This fails to capture the integrated hemodynamic phenotype of an individual. Consequently, it overlooks the significant heterogeneity in adaptive responses. This gap hinders the development of early, targeted risk stratification strategies. Notably, while emerging data-driven methods like unsupervised machine learning have proven effective in revealing clinically distinct subgroups in other cardiovascular populations^7^, and while the assessment of multi-system vascular impairment has been undertaken^2^, a systematic, phenotype-based investigation within healthy high-altitude dwellers remains lacking.

Recent advances in data-driven methodologies offer a path to address this limitation. In cardiovascular medicine, unsupervised machine learning techniques are increasingly applied to discover novel, clinically meaningful subgroups within seemingly uniform patient populations, revealing patterns not apparent through traditional analysis ^8,9^. Translating this paradigm to high-altitude physiology holds great promise. We hypothesize that clinically healthy individuals residing at high altitude naturally cluster into distinct hemodynamic adaptation phenotypes, and that these phenotypes are associated with a gradient of risk for multi-system subclinical target organ damage. To test this hypothesis, we conducted a cross-sectional study among a healthy high-altitude resident population. We employed an unsupervised machine learning (K-means clustering) approach on a comprehensive panel of hemodynamic variables to identify intrinsic adaptation phenotypes. Subsequently, we rigorously evaluated the association of these empirically derived phenotypes with a suite of validated indicators of vascular, cardiac, and renal health.

The primary objectives of this study were: To identify distinct hemodynamic adaptation phenotypes using an unsupervised clustering algorithm. To compare the burden of subclinical target organ damage, assessed by carotid intima-media thickness, left ventricular mass index, estimated glomerular filtration rate, and pulse wave velocity, across the identified phenotypes. By moving beyond isolated metrics to define integrated phenotypes, this work aims to reframe our understanding of cardiovascular adaptation at high altitude from a binary state to a spectrum of risk. Identifying high-risk hemodynamic phenotypes within the healthy population could enable earlier, more precise monitoring and pave the way for targeted interventions to prevent the progression to overt cardiovascular disease in high-altitude communities.

## 2 Methods

### 2.1 Study Population and Data Source

This cross-sectional study utilized data from a routine health examination cohort of adults residing in a high-altitude region (average altitude: 3,300 meters). Consecutive participants who underwent comprehensive cardiovascular assessments between October 2023 and October 2025 were considered for inclusion. Inclusion criteria were: (1) age ≥18 years; (2) permanent residency at high altitude (≥3,300 m) for more than one year; (3) availability of complete hemodynamic and target organ assessment data. Exclusion criteria included: (1) a known history of clinical cardiovascular disease (coronary artery disease, heart failure, stroke), chronic kidney disease (stage ≥3), or diabetes mellitus; (2) use of antihypertensive, lipid-lowering, or antidiabetic medications; (3) pregnancy; (4) presence of acute illness at the time of examination. According to the above criteria, 694 participants were finally included. The study protocol was reviewed and approved by the Institutional Review Board of Ruijin Hospital Ethics Committee, Shanghai Jiaotong University School of Medicine (Approval No. 2023-127, Registration number: ChiCTR2000040308). Of the 694 participants included in the final analysis, complete data for LVMI and PWV were available for 693 and 674 participants, respectively. Data for IMT and eGFR were available for 249 and 539 participants, respectively, due to the phased introduction of carotid ultrasound and serum creatinine measurements during the later stage of the study period.

### 2.2 Measurement of Hemodynamic and Target Organ Parameters

All measurements were performed by trained personnel following standardized protocols in a temperature-controlled room after participants had rested for at least 10 minutes.

#### 2.2.1 Hemodynamic Indicators

##### Peripheral and Central Blood Pressures

Peripheral systolic and diastolic blood pressures (pSBP, pDBP) were measured on the brachial artery using a validated automated oscillometric device Omron HEM-7206. Central systolic blood pressure (cSBP) and augmentation index corrected for a heart rate of 75 beats/min (AIx@75) were derived non-invasively from the radial artery pulse waveform obtained by applanation tonometry (SphygmoCor-XCEL).

##### Derived Parameters

Peripheral and central pulse pressures (pPP, cPP) were calculated as pSBP-pDBP and cSBP-cDBP, respectively. The peripheral-to-central pulse pressure ratio (pPP/cPP) and the peripheral-central systolic pressure difference (pSBP-cSBP) were also computed.

#### 2.2.2 Target Organ Damage Indicators

##### Vascular Structure

Carotid intima-media thickness (IMT) was measured on the far wall of the common carotid artery using high-resolution B-mode ultrasonography (Philips EPIQ 7]). The average of three measurements on each side was used for analysis.

##### Cardiac Structure

Left ventricular mass (LVM) was calculated from two-dimensional echocardiographic measurements according to the American Society of Echocardiography recommendations. LVM was indexed to body surface area to obtain the left ventricular mass index (LVMI).

##### Renal Function

Serum creatinine was measured using an enzymatic method. The estimated glomerular filtration rate (eGFR) was calculated using the Chronic Kidney Disease Epidemiology Collaboration (CKD-EPI) equation.

##### Arterial Stiffness

Carotid-femoral pulse wave velocity (PWV) was measured as the gold-standard index of arterial stiffness using a dedicated device (SphygmoCor-XCEL) by recording pulse waves at the carotid and femoral arteries.

### 2.3 Statistical Analysis

The statistical analysis followed a pre-defined sequential workflow. First, data preprocessing involved assessing normality for all variables using the Shapiro-Wilk test. Continuous variables are presented as mean ± standard deviation (SD) for normally distributed data or median (interquartile range) otherwise, while categorical variables are presented as frequencies (percentages); no imputation was performed for missing data. Subsequently, unsupervised K-means cluster analysis was employed to identify distinct hemodynamic phenotypes. The clustering variables—pSBP, cSBP, pPP, cPP, AIx@75, pPP/cPP, and pSBP-cSBP—were standardized (z-score transformation) prior to analysis. The optimal number of clusters (k=3) was determined by evaluating the elbow method (within-cluster sum of squares), silhouette score, and clinical interpretability using the stats package in R (version 4.5.2). Finally, differences across phenotypes were tested using one-way ANOVA (continuous variables) or chi-square tests (categorical variables). For significant ANOVA results, Bonferroni-corrected post-hoc pairwise comparisons were conducted, with a two-sided p-value < 0.05 after correction considered statistically significant.

### 2.4 Graphical Presentation

For graphical presentation, a grouped bar chart was generated to visualize the mean values of the four key target organ indicators (carotid intima-media thickness [IMT], estimated glomerular filtration rate [eGFR], left ventricular mass index [LVMI], and pulse wave velocity [PWV]) across the three hemodynamic phenotypes, with error bars representing the standard error of the mean (SEM) and statistical significance from Bonferroni-corrected post-hoc tests annotated (e.g., *, **, ***). Additionally, a conceptual hemodynamic adaptation risk spectrum diagram was created to illustrate the continuum of cardiovascular risk, positioning the three phenotypes along a gradient axis by integrating their defining hemodynamic profiles with the associated gradient of target organ damage burden. All statistical analyses were performed using R (version 4.5.2) or SPSS (version 29.0.2, IBM Corp.).

## 3 Results

### 3.1 Identification of Three Hemodynamic Phenotypes by Cluster Analysis

Unsupervised K-means cluster analysis was conducted on nine key hemodynamic variables—peripheral and central systolic blood pressure (pSBP, cSBP), peripheral and central pulse pressure (pPP, cPP), augmentation pressure (AP), augmentation index (AIx), heart rate-adjusted augmentation index (AIx@HR75), pulse pressure amplification ratio (pPP/cPP), and systolic pressure amplification (pSBP-cSBP)—which optimally partitioned the study population into three distinct phenotypes based on the elbow method and clinical interpretability (Figure 1, Table 1). These were defined as follows: Cluster 2 (C2, n=245): the “Balanced Adaptation” phenotype, exhibiting the most favorable hemodynamic profile with the lowest pSBP, cSBP, AP, AIx, and AIx@HR75, and the highest pPP/cPP; Cluster 1 (C1, n=267): the “Vascular Stress” phenotype, characterized by peripheral systolic pressure (125.9 ± 11.3 mmHg) comparable to C2 but with significantly elevated wave reflection indices, demonstrating the highest AIx (29.7 ± 9.2%) and AIx@HR75 (31.9 ± 9.7%) and the lowest pPP/cPP; and Cluster 3 (C3, n=182): the “High-Load Decompensation” phenotype, displaying markedly elevated hemodynamic load with the highest values for pSBP, cSBP, pPP, and cPP.

**Figure 1.**
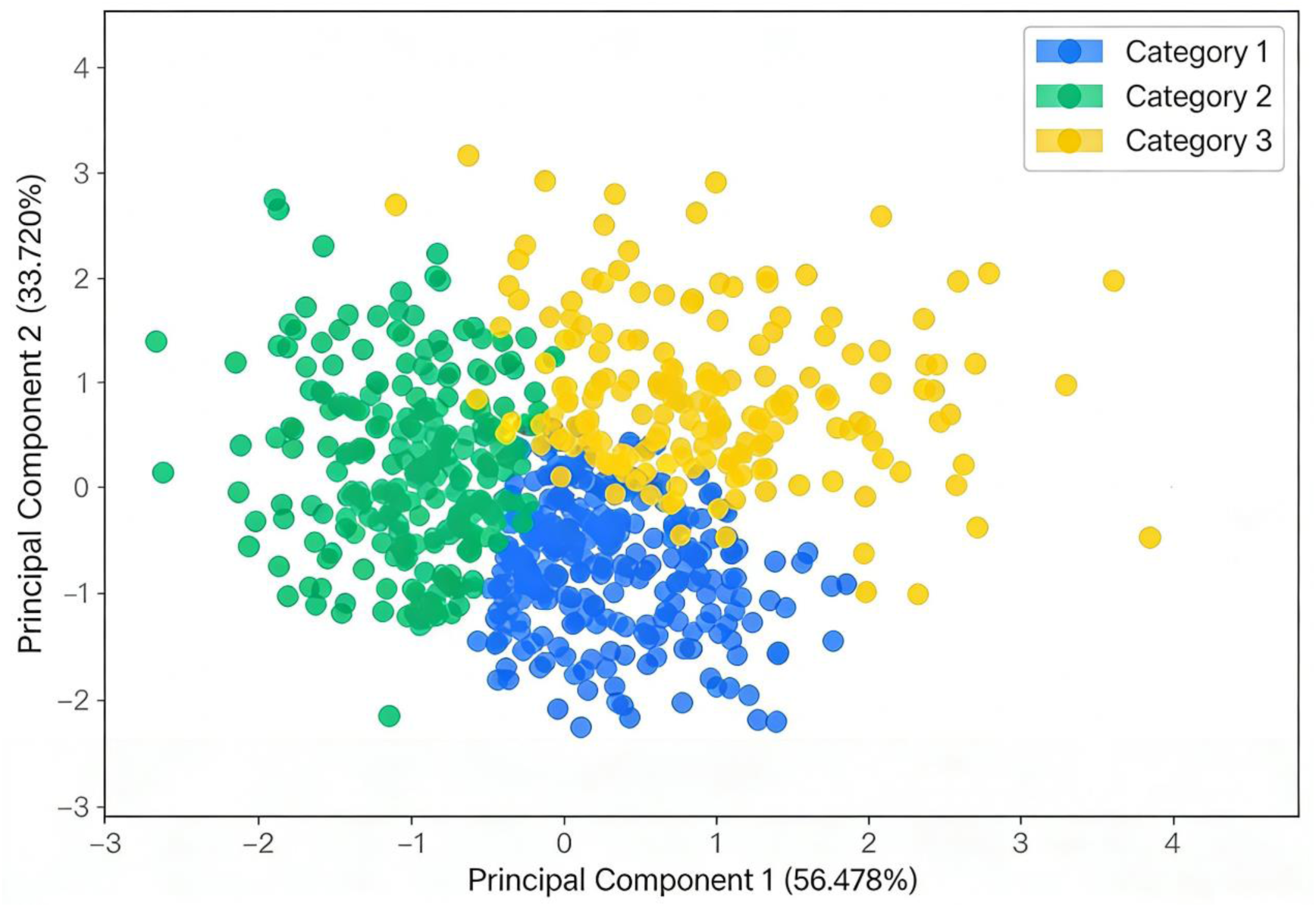
Scatter plot of the three hemodynamic phenotypes based on principal component analysis. * Scatter plot of the study population projected onto the first two principal components (PC1 and PC2) derived from nine hemodynamic variables (peripheral and central systolic blood pressure, peripheral and central pulse pressure, augmentation pressure, augmentation index, heart rate-adjusted augmentation index, peripheral-to-central pulse pressure ratio, and systolic pressure amplification). PC1 explained 56.478% of the total variance, and PC2 explained 33.720% of the total variance. Each point represents an individual participant, colored according to their assigned phenotype: C1 (Vascular Stress, blue), C2 (Balanced Adaptation, green), and C3 (High-Load Decompensation, red). The clear spatial separation of the three clusters validates the robustness of the three-cluster solution identified by K-means clustering. *

**Table 1.**
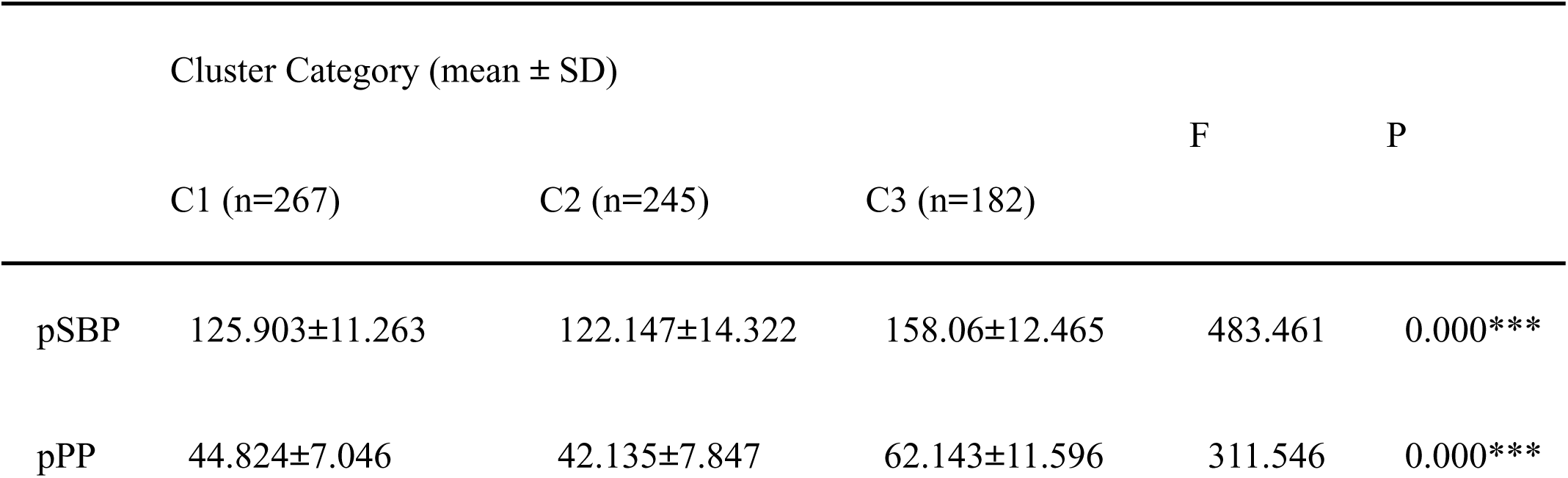
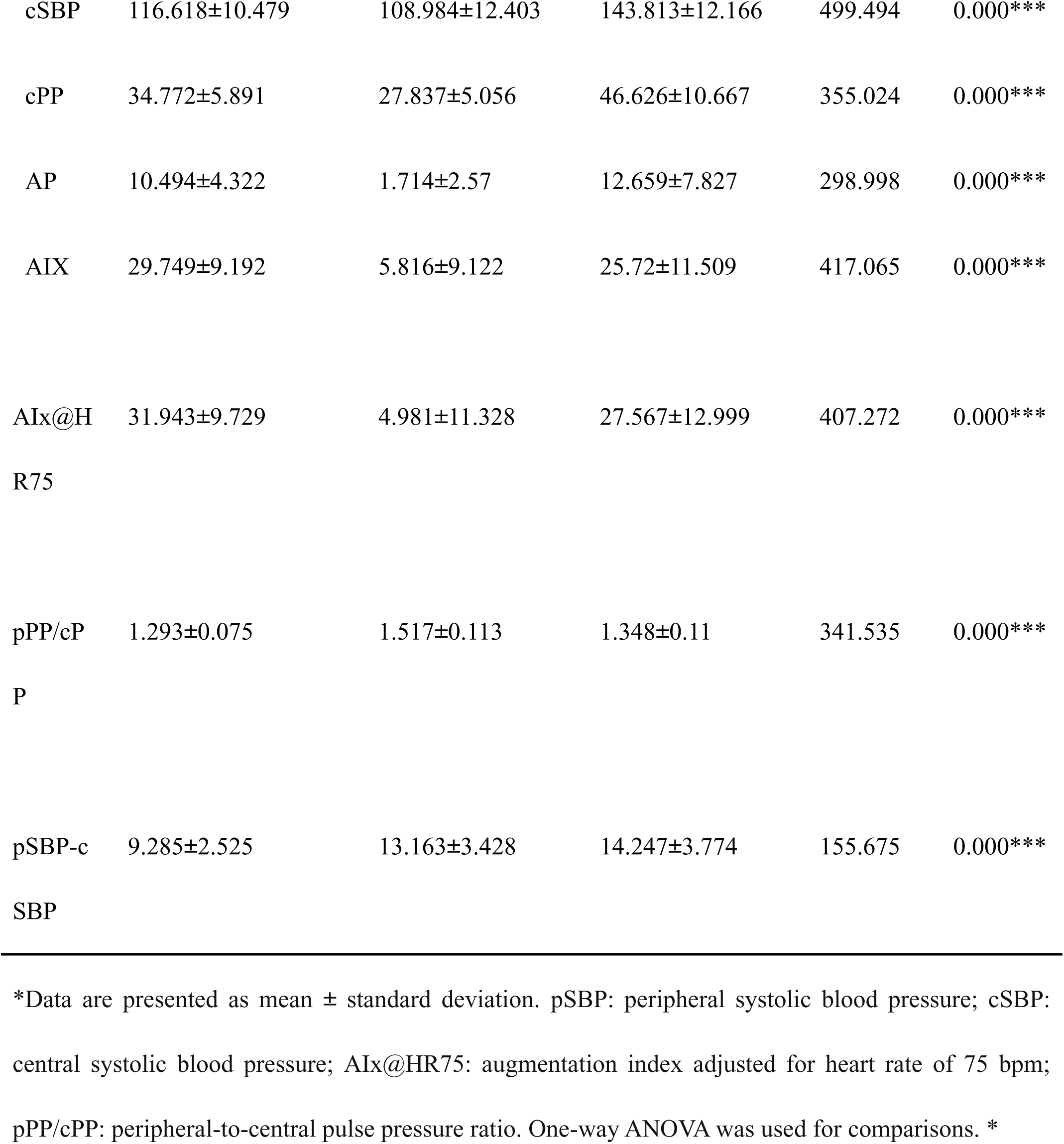
Hemodynamic Characteristics of the Three Identified Phenotypes (Cluster Solution)

### 3.2 Baseline Characteristics Across Phenotypes

The baseline demographic and clinical characteristics of the total cohort and the three phenotypes are summarized in Table 2. Significant differences were observed among the phenotypes in age, sex distribution, smoking status, and heart rate (all P < 0.01).

**Table 2.** Baseline Characteristics of the Study Population and Phenotypes.

| Characteristics | C3 | C2 | C1 | P value |
| --- | --- | --- | --- | --- |
| n | 182 | 245 | 267 |  |
| SEX, n (%) |  |  |  | < 0.001 |
| Female | 77 (11.1%) | 107 (15.4%) | 162 (23.3%) |  |
| Male | 105 (15.1%) | 138 (19.9%) | 105 (15.1%) |  |
| AGE, median (IQR) | 58.5 (50, 65) | 45 (36, 55) | 55 (48, 63) | < 0.001 |
| BMI, median (IQR) | 25.67 (23.62, 28.82) | 25.26 (22.25, 28.91) | 25.07 (22.96, 27.55) | 0.075 |
| SMOKE, n (%) |  |  |  | 0.002 |
| No | 144 (20.8%) | 176 (25.4%) | 226 (32.6%) |  |
| Yes | 37 (5.3%) | 69 (10%) | 41 (5.9%) |  |
| HbA1c, median (IQR) | 5.9 (5.475, 6.725) | 5.8 (5.4, 6.9) | 5.9 (5.6, 6.6) | 0.551 |
| IMT, median (IQR) | 1.2 (1, 1.3) | 0.95 (0.8, 1.2) | 1.1 (0.9, 1.2) | < 0.001 |
| EGFR, median (IQR) | 96 (80.25, 104) | 99.5 (87, 109) | 97 (87, 106) | 0.010 |
| LVMI, mean ± SD | 69.18 ± 40.73 | 48.19 ± 45.64 | 60.43 ± 44.65 | < 0.001 |
| Average PWV, median | 9.1 (8, 10.3) | 7.2 (6, 8.2) | 7.6 (6.6, 8.9) | < 0.001 |
| (IQR) |  |  |  |  |
| HR, mean ± SD | 70.264 ± 11.496 | 77.143 ± 13.004 | 69.375 ± 11.025 | < 0.001 |
| pSBP, median (IQR) | 155 (149, 166) | 122 (112, 134) | 128 (118, 135) | < 0.001 |
| pPP, median (IQR) | 61 (54, 68) | 42 (36, 47) | 44 (40, 50) | < 0.001 |
| cSBP, median (IQR) | 142 (135, 152) | 109 (100, 118) | 117 (109, 124) | < 0.001 |
| cPP, median (IQR) | 45 (39, 52) | 28 (24, 31) | 34 (30, 38.5) | < 0.001 |
| AP, median (IQR) | 11.5 (7, 16) | 2 (0, 4) | 10 (7, 13) | < 0.001 |
| AIX, median (IQR) | 26 (18, 32) | 7 (1, 13) | 28 (22, 35) | < 0.001 |
| AIX@HR75, median (IQR) | 27.9 (19.32, 36.053) | 6.22 (-2.29, 14.27) | 29.71 (24.885, 39.145) | < 0.001 |
| pPP/cPP, median (IQR) | 1.33 (1.28, 1.4175) | 1.5 (1.43, 1.58) | 1.3 (1.24, 1.345) | < 0.001 |
| pSBP-cSBP, median (IQR) | 14 (12, 16) | 13 (11, 15) | 9 (7.5, 11) | < 0.001 |

\*Variables include age, sex, smoking status, heart rate, and baseline subclinical target organ damage indicators (carotid IMT, eGFR, LVMI, PWV). Significant differences among phenotypes were observed for age, sex, smoking status, heart rate, IMT, eGFR,
LVMI, and PWV (all $P < 0.05$ ). C2 (Balanced Adaptation) was the youngest, whereas C3 (High-Load Decompensation) was the oldest\*

The C2 (Balanced Adaptation) phenotype was the youngest, while the C3 (High-Load Decompensation) phenotype was the oldest. Notably, despite these baseline differences, the key indicators of subclinical target organ damage—carotid intima-media thickness (IMT), estimated glomerular filtration rate (eGFR), left ventricular mass index (LVMI), and pulse wave velocity (PWV)—already showed significant inter-group variations at baseline (all P < 0.05).

### 3.3 Gradient of Subclinical Target Organ Damage

The burden of subclinical damage across the four organ systems exhibited a clear gradient associated with the hemodynamic phenotypes (Table 3). One-way ANOVA revealed highly significant overall differences among the three groups for IMT, eGFR, LVMI, and PWV (all P < 0.01).

**Table 3.**
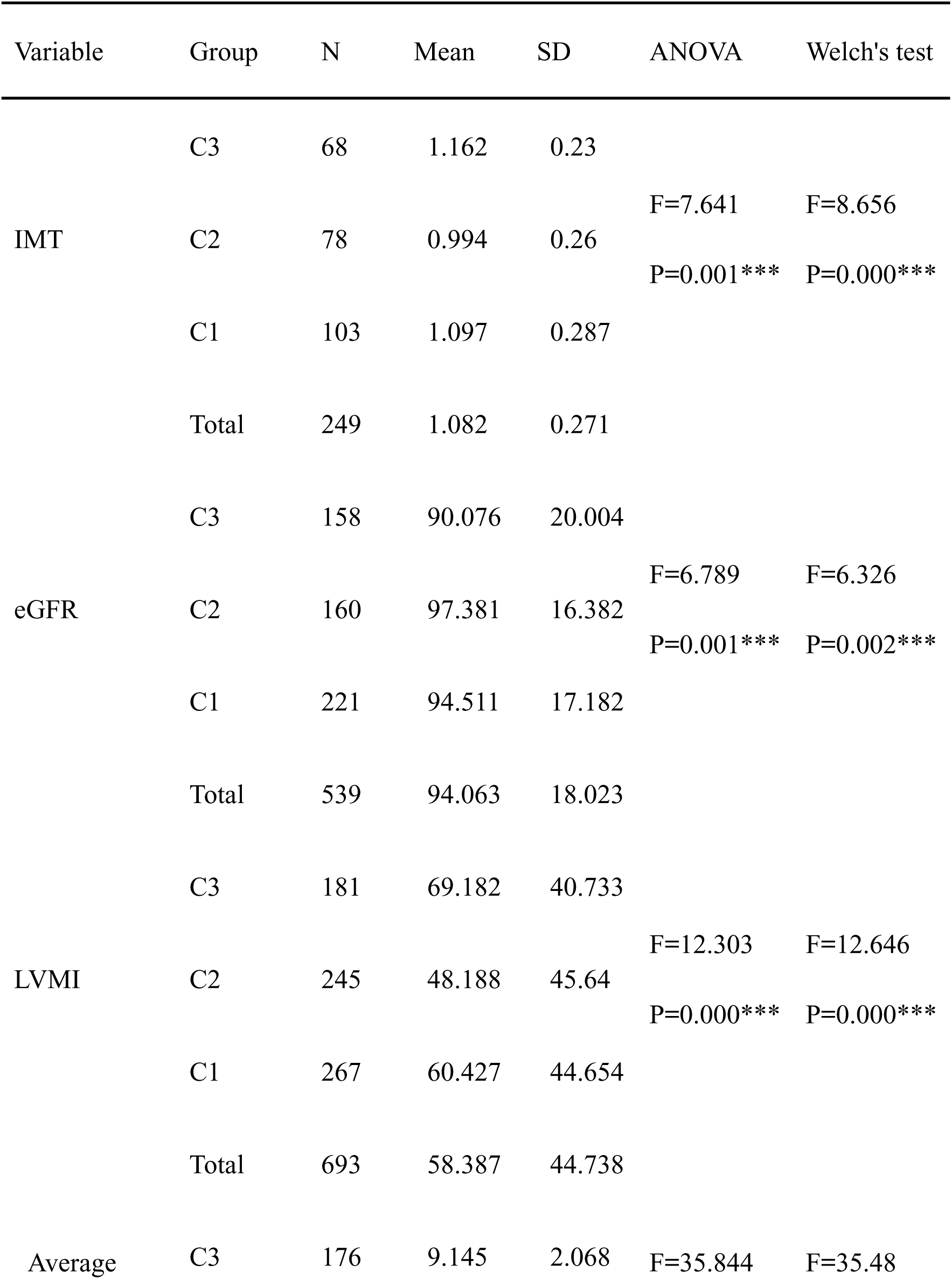
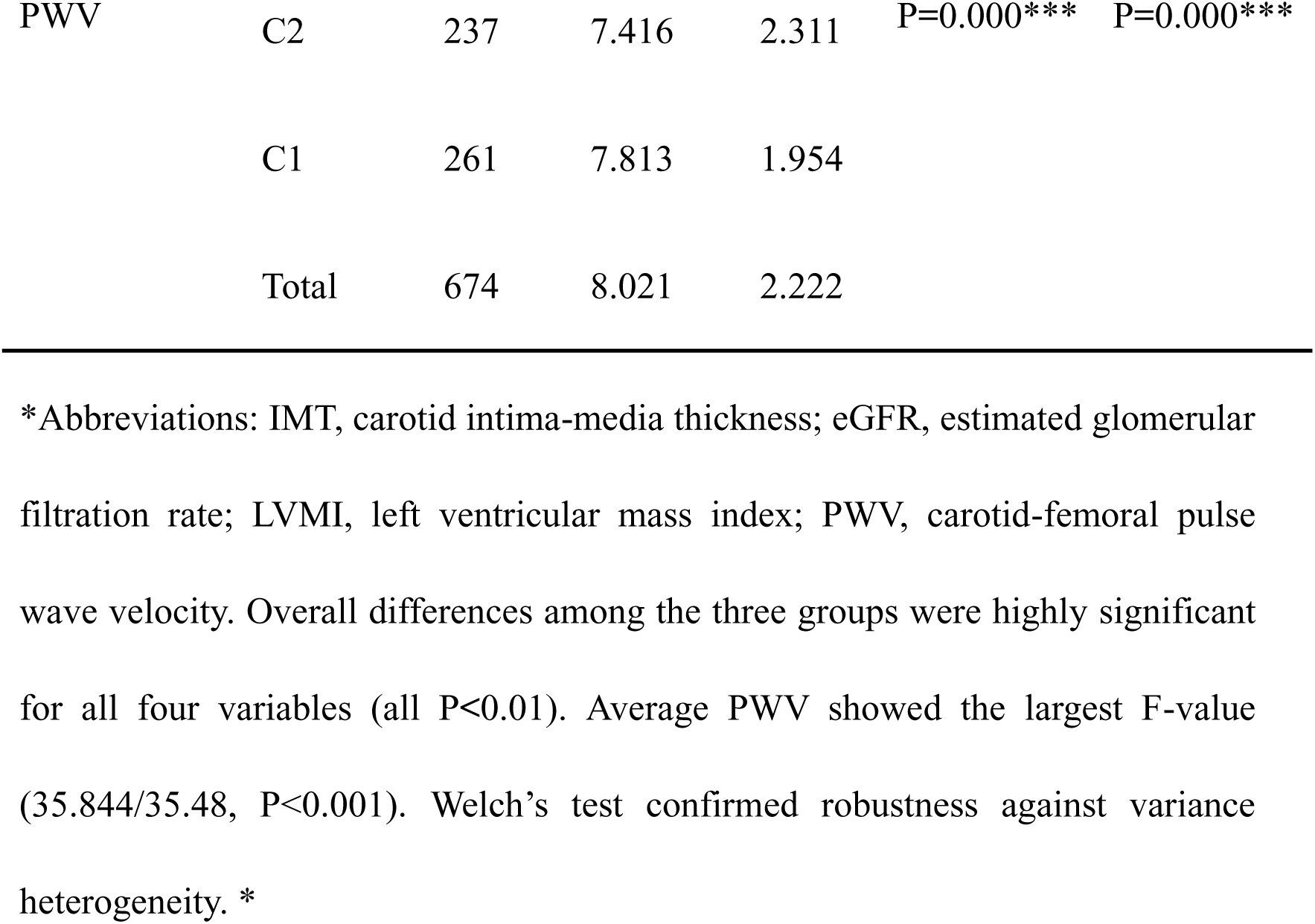
Analysis of Target Organ Damage Indicators Across Phenotypes.

Specifically, for IMT, the ANOVA (F = 7.641, P = 0.001) and Welch‘s test (F = 8.656, P < 0.001) consistently showed significant differences among the three groups. The C2 phenotype had the lowest mean IMT (0.994 ± 0.26 mm), while the C3 phenotype had the highest (1.162 ± 0.23 mm). For eGFR, significant differences were also observed (ANOVA: F = 6.789, P = 0.001; Welch’s F = 6.326, P = 0.002), with the C2 phenotype showing a significantly higher mean eGFR (97.38 ± 16.38 mL/min/1.73 m²) compared with the other two groups (C1: 94.51 ± 17.18; C3: 90.08 ± 20.00). LVMI demonstrated the most pronounced intergroup differences (ANOVA: F = 12.303, P < 0.001; Welch‘s F = 12.646, P < 0.001), with the lowest mean value in the C2 phenotype (48.19 ± 45.64 g/m²) and the highest in the C3 phenotype (69.18 ± 40.73 g/m²). PWV showed the largest F-value (ANOVA: F = 35.844, P < 0.001; Welch’s F = 35.48, P < 0.001), with the C3 phenotype exhibiting a markedly higher mean PWV (9.15 ± 2.07 m/s) compared with the C2 (7.42 ± 2.31 m/s) and C1 (7.81 ± 1.95 m/s) phenotypes.

Post-hoc pairwise comparisons with Bonferroni correction detailed the specific inter-phenotype differences (Table 4). For IMT, the mean difference between C1 and C2 was 0.103 mm (P = 0.010), and between C2 and C3 was −0.168 mm (P < 0.001); however, the difference between C1 and C3 did not reach statistical significance (P = 0.118). For eGFR, the largest mean difference was observed between C2 and C3 (7.31 mL/min/1.73 m², P < 0.001), and a significant difference was also found between C1 and C3 (4.44 mL/min/1.73 m², P = 0.017), whereas C1 and C2 were not significantly different (P = 0.122). All pairwise comparisons for LVMI were significant: C1 vs. C2 (12.24 g/m², P = 0.002), C2 vs. C3 (−20.99 g/m², P < 0.001), and C1 vs. C3 (−8.76 g/m², P = 0.039). For PWV, all pairwise comparisons were also significant: C1 vs. C2 (0.40 m/s, P = 0.037), C1 vs. C3 (−1.33 m/s, P < 0.001), and C2 vs. C3 (−1.73 m/s, P < 0.001).

**Table 4.**
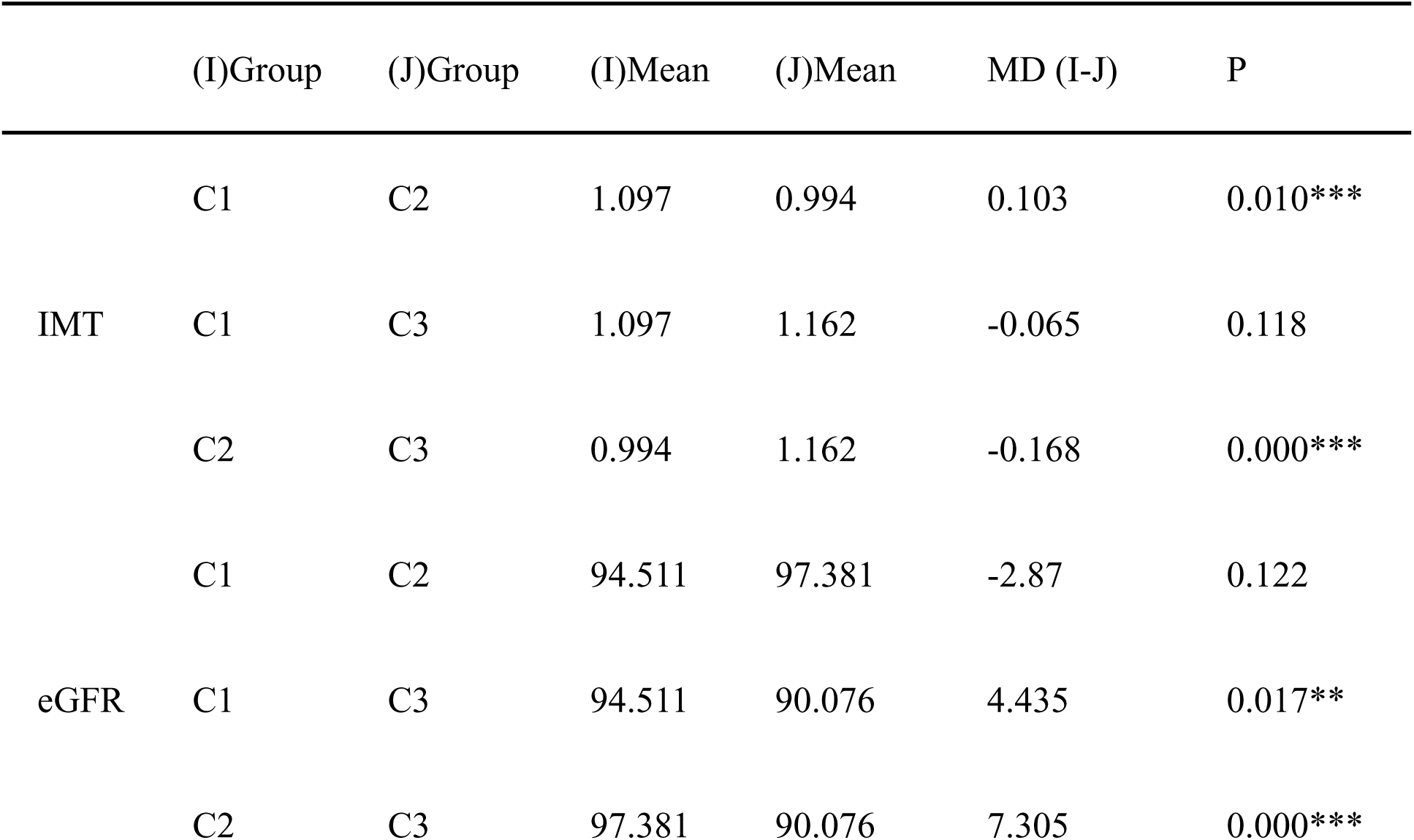
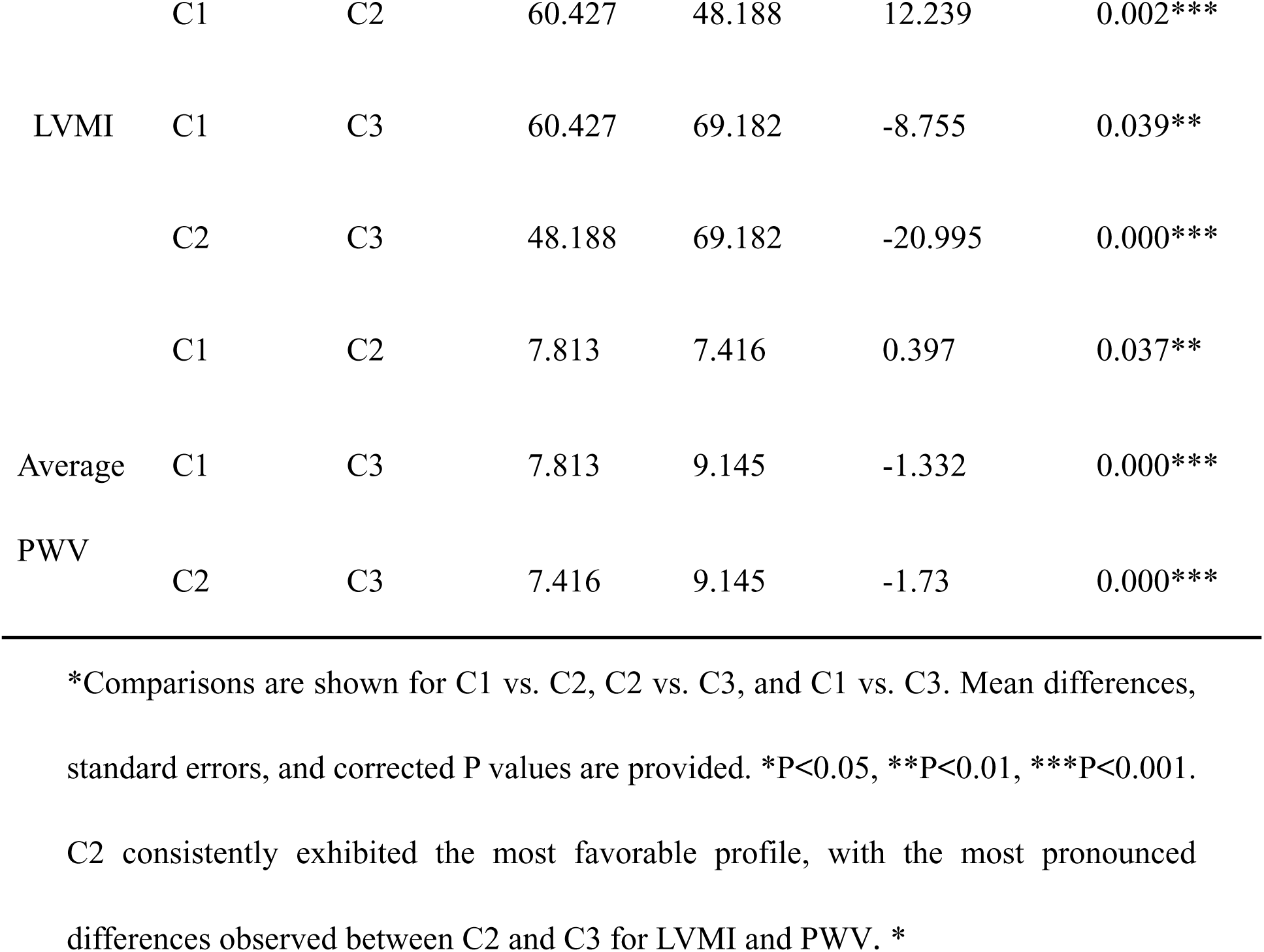
Post-Hoc Pairwise Comparisons (Bonferroni-adjusted) of Target Organ Damage.

Bar graphs (Figure 2) display the mean values of (A) IMT, (B) eGFR, (C) LVMI, and (D) PWV for the three phenotypes. Error bars represent the standard error of the mean (SEM). Significant differences between groups, determined by one-way ANOVA with Bonferroni-corrected post-hoc tests, are indicated as *P < 0.05, **P < 0.01, ***P < 0.001. The C2 phenotype consistently shows the most favorable values across all four indicators. A clear risk gradient is evident from C2 (lowest risk) to C1 (intermediate risk) to C3 (highest risk), particularly for LVMI and PWV.

**Figure 2.**
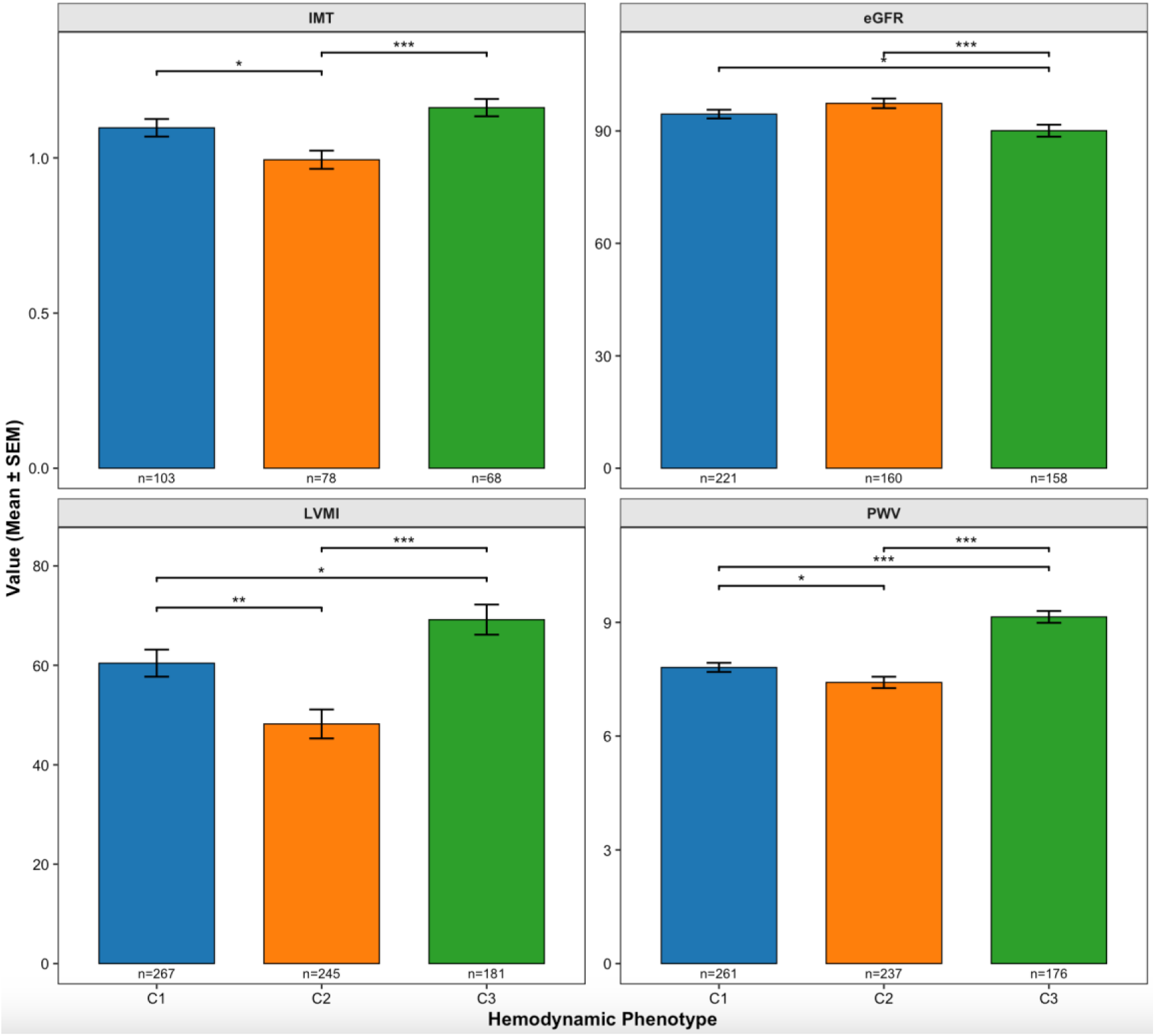
Comparison of target organ indicators across three hemodynamic phenotypes in high-altitude healthy population. *IMT (carotid intima-media thickness), eGFR (estimated glomerular filtration rate), LVMI (left ventricular mass index), PWV (pulse wave velocity). Phenotypes: C1 (Vascular Stress, blue), C2 (Balanced Adaptation, orange), C3 (High-Load Decompensation, green). Significance codes: *p<0.05, **p<0.01, ***p<0.001 (Bonferroni-corrected post-hoc tests). *

### 3.4 Conceptual Hemodynamic Adaptation Risk Spectrum

Synthesizing the distinct hemodynamic profiles and the graded burden of multi-organ damage, we propose a conceptual “Hemodynamic Adaptation Risk Spectrum” (Figure 3). This model positions the C2 (Balanced Adaptation) phenotype at the optimal, low-risk end, characterized by efficient hemodynamics and minimal subclinical injury. The C1 (Vascular Stress) phenotype occupies a crucial intermediate position, where normal-range peripheral blood pressure masks significant vascular stiffening and early intermediate organ damage, representing a likely pre-clinical high-risk state. The C3 (High-Load Decompensation) phenotype anchors the high-risk end, defined by overt pressure overload and the most severe multi-organ impairment.

**Figure 3.**
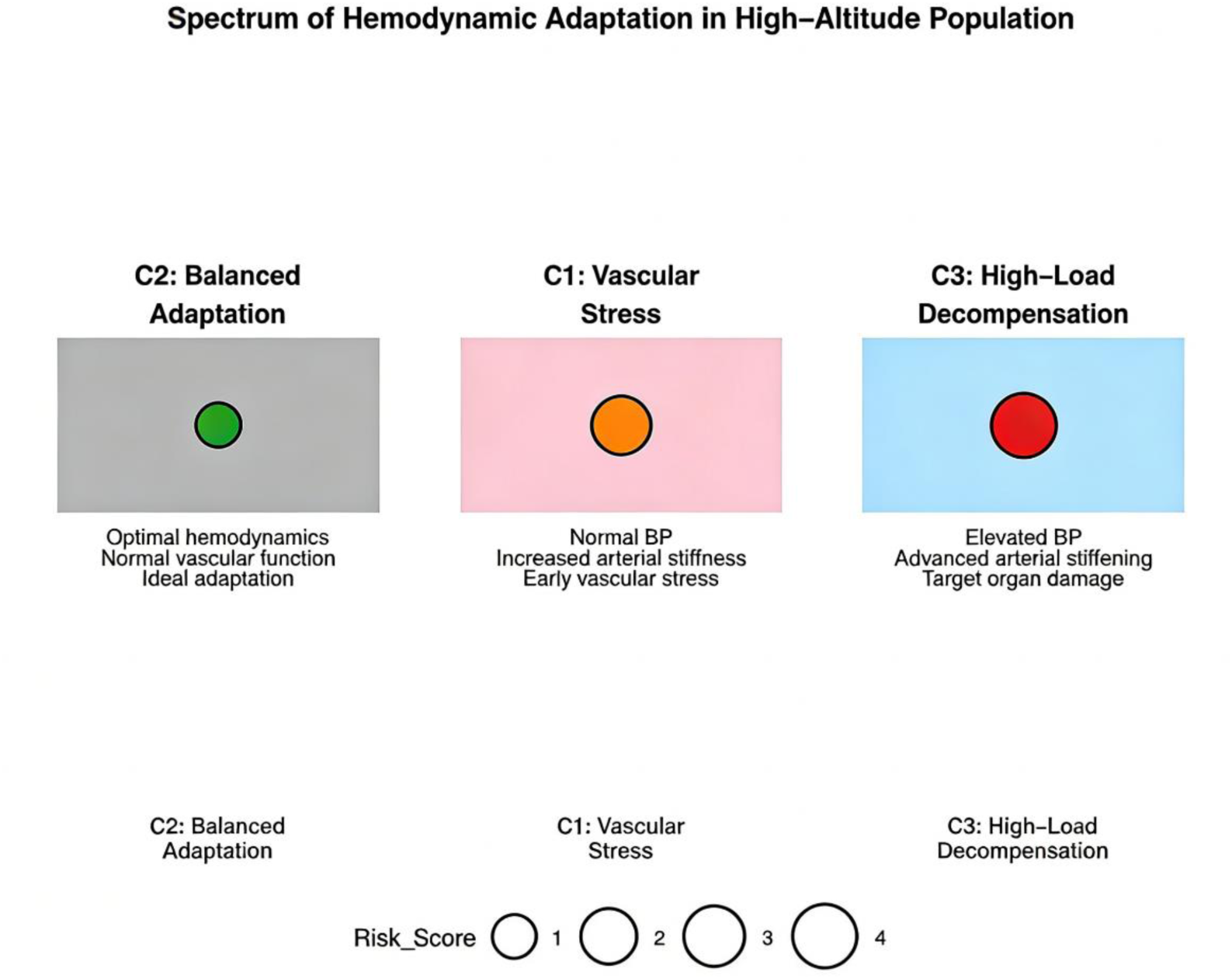
Spectrum of Hemodynamic Adaptation in High-Altitude Population. *Conceptual Hemodynamic Adaptation Risk Spectrum. C2 (Balanced Adaptation) is positioned at the low-risk end, characterized by efficient hemodynamics and minimal subclinical organ damage. C1 (Vascular Stress) occupies an intermediate, pre-clinical high-risk state in which normal-range peripheral blood pressure masks vascular stiffening and early target organ damage. C3 (High-Load Decompensation) anchors the high-risk end with overt pressure overload and severe multi-organ impairment. *

## 4 Discussion

### 4.1 Principal Findings

This study provides novel evidence for the heterogeneity of hemodynamic adaptation among healthy high-altitude dwellers, extending observations of a functional vascular spectrum within this population^2^. Using an unsupervised machine learning approach, we identified and validated three distinct hemodynamic phenotypes: Balanced Adaptation (C2), Vascular Stress (C1), and High-Load Decompensation (C3). These phenotypes were not arbitrarily defined but emerged naturally from the data, forming a coherent risk spectrum. Crucially, this spectrum was strongly associated with a gradient of multi-system subclinical target organ damage^10^.The most significant finding is the identification of the Vascular Stress (C1) phenotype, characterized by normal peripheral blood pressure coupled with significantly elevated arterial wave reflection, which exhibited an intermediate risk profile for early organ injury.

### 4.2 Interpretation of the “Vascular Stress” Phenotype and the Risk Spectrum

The Vascular Stress (C1) phenotype represents a critical intermediate state in the adaptation continuum. Its defining feature—preserved brachial systolic pressure but markedly increased augmentation index (AIx@75) and reduced pulse pressure amplification (pPP/cPP)—suggests a pathophysiological shift where increased arterial stiffness and enhanced wave reflection precede the development of overt hypertension, a hemodynamic consequence well-explained by the principles of wave reflection in stiffened arteries^11^. This pattern is consistent with the known effects of chronic hypoxia on vascular function, promoting endothelial dysfunction and sympathetic activation^12^ and involving maladaptive molecular pathways such as the HIF signaling axis, which augment pressure wave reflections from the periphery^13^. The sustained enhancement of peripheral chemoreceptor (e.g., carotid body) tonic activity under chronic hypoxia may underpin this persistent sympathetic and vascular tone^14^. Our finding that this phenotype already exhibits a greater burden of carotid thickening and cardiac remodeling than the Balanced Adaptation (C2) phenotype underscores its clinical relevance. It challenges the sole reliance on peripheral blood pressure for risk assessment at high altitude and highlights central hemodynamics and arterial stiffness as earlier, more sensitive markers of cardiovascular maladaptation, a notion robustly supported by evidence linking arterial stiffness to cardiovascular risk independent of blood pressure levels^15^.

The conceptual Hemodynamic Adaptation Risk Spectrum integrating C2, C1, and C3 provides a paradigm shift from a binary view of adaptation. It posits that cardiovascular risk at high altitude exists along a continuum, with the Vascular Stress phenotype occupying the pivotal transition zone from optimal compensation to decompensation. This model offers a framework for understanding the progressive nature of altitude-related vascular stress and identifies C1 as a potential target for pre-emptive intervention to halt progression towards overt disease.

### 4.3 Comparison with Previous Studies and Potential Mechanisms

Previous research has established that high-altitude residence is associated with higher prevalence of hypertension and subclinical vascular changes on a population level^16,17^. Our findings extend this knowledge by demonstrating that within a nominally healthy population, distinct adaptive pathways exist, each carrying a different risk. The High-Load Decompensation (C3) phenotype aligns with classical descriptions of high-altitude hypertension and its associated cardiac and vascular damage. The novel Vascular Stress (C1) phenotype, however, may explain why some individuals with “normal” blood pressure still develop premature cardiovascular issues in high-altitude regions.

The physiological basis for these phenotypes likely involves differential activation of adaptive and maladaptive pathways to chronic hypoxia. The favorable profile of the C2 phenotype may reflect optimal balance in sympathetic tone, nitric oxide bioavailability, and vascular compliance^18^. In contrast, the C1 phenotype suggests a predominant maladaptive response in the vascular wall—increased stiffness and wave reflection—possibly driven by hypoxia-induced endothelial dysfunction, oxidative stress, and altered collagen metabolism, without a concomitant strong pressor response. The C3 phenotype likely represents a further decompensated state where neurohormonal activation (e.g., RAAS, sympathetic system) supervenes, leading to sustained pressure overload^19^.

### 4.4 Clinical and Public Health Implications

Our phenotype-based stratification has direct implications for advancing precision prevention strategies in high-altitude communities. First, it argues for a paradigm shift in screening protocols. Moving beyond the sole reliance on brachial blood pressure, routine health assessments should incorporate evaluations of arterial stiffness (e.g., via pulse wave analysis for augmentation index or carotid-femoral pulse wave velocity) to specifically identify individuals with the Vascular Stress (C1) profile, who are missed by conventional metrics^20^. Second, this approach enables tailored risk communication; individuals identified as C1 can be counseled on their elevated cardiovascular risk despite normal peripheral blood pressure, which may enhance motivation for lifestyle modifications^21^. From a public health perspective, this stratification allows for more efficient resource allocation, enabling programs to prioritize monitoring and management of the higher-risk C1 and C3 subgroups, thereby potentially improving the cost-effectiveness of cardiovascular prevention initiatives^22^. Most importantly, the C1 phenotype delineates a clear preclinical target for intervention trials. It presents a unique opportunity to test whether early, targeted interventions—such as specific exercise regimens, dietary nitrate supplementation, or novel vasodilators aimed at improving endothelial function and reducing arterial stiffness—can halt or reverse the progression to overt hypertension and organ damage^23^.

### 4.5 Strengths and Limitations

The main strengths of this study include the use of an unbiased, data-driven clustering method to define phenotypes^24^, the comprehensive multi-system evaluation of subclinical damage using standard techniques, and the integration of findings into a novel conceptual model.

This study also has limitations. First, its cross-sectional design precludes causal inferences about the relationship between phenotypes and organ damage, or knowledge of the longitudinal stability of these phenotypes. Second, although statistically significant, the observed differences in age and heart rate among phenotypes are potential confounding factors that residual confounding may influence the results. Future studies with longitudinal design and adjusted analyses are warranted^25^. Third, the sample was from a single region, and generalizability to other high-altitude populations with different genetic backgrounds (e.g., Andean, Ethiopian) requires validation^26^. Finally, the absence of data on potential biological mediators—such as sympathetic activity, specific inflammatory markers, or genetic polymorphisms—limits our ability to elucidate the underlying mechanisms responsible for the genesis of these distinct phenotypes^27^.

## 5 Conclusions and Future Perspectives

In conclusion, our study demonstrates that hemodynamic adaptation to high altitude in healthy individuals is not uniform but manifests as distinct phenotypes arrayed along a spectrum of cardiovascular risk. The identification of the Vascular Stress phenotype—a high-risk state masked by normal peripheral blood pressure—is a key finding that calls for a revision of current risk assessment strategies in high-altitude populations. The proposed Hemodynamic Adaptation Risk Spectrum provides a useful framework for understanding the progression of cardiovascular maladaptation. First, future research should conduct longitudinal cohort studies to track the temporal stability of these hemodynamic phenotypes and establish their prognostic value for predicting hard clinical endpoints such as cardiovascular events^28^. Subsequently, employing multi-omics approaches—including genomics, metabolomics, and proteomics—is essential to elucidate the underlying biological determinants and molecular pathways that distinguish each phenotype^29,30^. Building on this mechanistic understanding, the next critical step is to design and test targeted intervention trials, particularly for individuals within the Vascular Stress group, to evaluate strategies for halting or reversing disease progression. Ultimately, the systematic integration of such phenotypic profiling into high-altitude clinical practice could pave the way for more personalized and effective cardiovascular prevention, transforming risk stratification from a population-based to an individual-centered paradigm.

## Acknowledgments

The authors would like to thank all participants who volunteered for this study. We are grateful to the medical staff at Deqing Tibetan Autonomous Prefecture People’s Hospital for their assistance with data collection and participant management. All persons acknowledged have provided written consent for their inclusion in this section.

## Ethics of approval statement

The study protocol was reviewed and approved by the Ethics Committee of Ruijin Hospital, Shanghai Jiaotong University School of Medicine (2023-127).

